# Pre-meiotic, 24-nt reproductive phasiRNAs are abundant in anthers of wheat and barley but not rice and maize

**DOI:** 10.1101/2020.06.18.160440

**Authors:** Sébastien Bélanger, Suresh Pokhrel, Kirk Czymmek, Blake C. Meyers

## Abstract

Two classes of pre-meiotic (21-nt) and meiotic (24-nt) phasiRNAs and their patterns of accumulation have been described in maize and rice anthers. Their precise function remains unclear, but some studies have shown that they support male fertility. The important role of phasiRNAs in anthers underpins our current study to their characterization in wheat and barley anthers. In this study, we staged anthers at every 0.2 mm of development for one wheat and two barley varieties. We isolated pre-meiotic (0.2 mm, 0.4 mm and 0.6 mm), meiotic (0.8 mm, 1.0 mm and 1.4 mm) and post-meiotic (1.8 mm) anthers for which we then investigated accumulation patterns of RNAs, including reproductive phasiRNAs. We annotated a total 12,821 and 2,897 *PHAS* loci in the wheat and barley genomes, respectively. When comparing the total number of *PHAS* loci in genomes of maize, rice, barley and wheat, we characterized an expansion of reproductive *PHAS* loci in the genomes of *Poaceae* subfamilies from *Panicoideae* to *Oryzoideae* and to *Poideae*. In addition to the two classes of pre-meiotic (21-nt) and meiotic (24-nt) phasiRNAs, previously described in maize and rice anthers, we described a group of 24-nt phasiRNAs that accumulate in pre-meiotic anthers. The absence of pre-meiotic 24-nt phasiRNAs in maize and rice suggests a divergence in grass species of the *Poideae* subfamily. Additionally, we performed a co-expression gene analysis describing the regulation of phasiRNA biogenesis in wheat and barley anthers. We highlight *AGO9* and *AGO6* as candidate binding partners of pre-meiotic and meiotic 24-nt phasiRNAs, respectively.

**One sentence summary:** In wheat and barley anthers, 24-nt reproductive phasiRNAs are abundant in both meiotic and pre-meiotic stages.

## INTRODUCTION

Plants produce small RNAs (sRNAs) of typically 21 to 24 nucleotides (nt). The biogenesis and regulation of sRNAs requires RNA polymerases (Pol), Dicer-like (DCL) proteins, double-stranded RNA-binding (DRB) proteins, RNA-directed RNA polymerases (RDRs) and Argonaute (AGO) proteins (Borges and Martienssen, 2015; Yu et al., 2018). Among the sRNA expressed in plants, phased siRNAs, or “phasiRNAs”, are a distinct group of sRNAs characterized as a product of processive cleavage of double-stranded RNAs in regular increments (duplexes of 21 nt or 24 nt) from a well-defined terminus (Axtell and Meyers, 2018). PhasiRNA biogenesis initiates via microRNA-directed (miRNA), AGO-catalyzed cleavage of a single-stranded RNA precursor which is then converted to double-stranded RNA by an RDR protein, prior to being processed into 21- or 24-nt RNA duplexes by a DCL protein. PhasiRNAs originate from both protein-coding and non-coding transcripts (Fei et al., 2013; Komiya, 2017; Yu et al., 2018). PhasiRNAs originating from protein-coding genes are 21-nt long (Komiya, 2017; Yu et al., 2018). A particular group of phasiRNAs, called reproductive phasiRNAs, is specific to male reproductive tissue and can be either 21- or 24-nt long (Johnson et al., 2009; Song et al., 2012a; Zhai et al., 2015; Fei et al., 2016; Komiya, 2017; Yu et al., 2018). Accumulation of two distinct groups of pre-meiotic (21-nt) and meiotic (24-nt) reproductive phasiRNAs was found to occur in maize (*Zea mays*; Zhai et al., 2015) and rice (*Oryza sativa*; Komiya et al., 2014; Fei et al., 2016) anthers. Subsequent work has demonstrated the widespread prevalence of 24-nt phasiRNAs in angiosperm species (Xia et al., 2019). While pathways governing reproductive phasiRNA production have been described, there remain gaps in our understanding.

Nonomura et al. (2007) identified the *MEIOSIS ARRESTED AT LEPTOTENE 1* (*MEL1*) gene, which encodes an AGO protein, in a male-sterile rice mutant. A *mel1* loss-of-function mutant is characterized by an arrest of chromosome condensation at early meiotic stages as well as by irregularly sized, multi-nucleated and vacuolated pollen mother cells (Nonomura et al., 2007). Komiya et al. (2014) demonstrated that MEL1, also known as OsAGO5c, is a binding partner specific to 21-nt phasiRNAs that are generated from long non-coding RNAs (lncRNAs). Active in pre-meiotic male reproductive tissues, production of these 21-nt phasiRNAs is dependent on miR2118, *RDR6, DCL4, MEL1*, and presumably a copy of AGO1 to load miR2118 (Song et al., 2012a; Song et al., 2012b; Komiya et al., 2014; Yu et al., 2018). Although most of the 21-nt phasiRNA loci are associated with male reproductive development, 21-nt phasiRNAs are also involved in other biological processes (Fei et al., 2013). For instance, the cleavage of *TAS3* precursor transcripts triggered by miR390 is required for normal development in most if not all land plants (Xia et al., 2017). In both cases, RDR6 and DCL4 proteins process the 21-nt phasiRNAs, which complicates the study of this pathway with respect to reproductive development.

A second size class of phasiRNAs, 24-nt phasiRNAs, is apparently specific to male reproductive development in angiosperm species. Although our understanding of this pathway and its function is incomplete, several studies have begun to provide details. First, we know that 24-nt phasiRNA production is dependent on miR2275, *RDR6, DCL5*, and presumably a copy of AGO1 to load miR2275 (Kapoor et al., 2008; Song et al., 2012a; Song et al., 2012b; Yu et al., 2018). *DCL5* is a monocot-specific, *Dicer-like* gene, that is exclusive to the 24-nt phasiRNA pathway. Recently, a *dcl5* loss-of-function mutation was shown to result in developmental defects in maize tapetum (Teng et al., 2020). Although it is not yet confirmed, evidence based on transcriptional regulation suggests that AGO2b (Fei et al., 2016) and AGO18 (Zhai et al., 2015; Fei et al., 2016) might be the binding partners of 24-nt phasiRNA (analogous to MEL1, for 21-nt phasiRNAs.)

The precise function of reproductive phasiRNAs remains unknown, but some studies suggest that they may play a critical role in maintaining fertility under environmental stresses. For example, a mutation in the rice *PMS1T* transcript (a lncRNA that is a 21-nt phasiRNA precursor, or *PHAS* transcript) results in photoperiod-sensitive genic male sterility (Fan et al., 2016). Furthermore, the phenotype of temperature-sensitive male sterility was observed in a maize *dcl5* loss-of-function mutant (Teng et al., 2020). Thus, both, 21-nt and 24-nt phasiRNA pathways appear to be vitally involved in reproductive biology and may provide approaches to utilize self-fertilized grass species in hybrid seed production. Bread wheat (*Triticum aestivum*) and barley (*Hordeum vulgare ssp. vulgare*) are two self-fertilized species that respectively rank 1st and 4th among economically important cereal crops (http://www.fao.org/faostat/). A full understanding of the gene regulatory networks controlling anther development is crucial as it may enable regulation of pollen production and the development of hybrid seed technologies for plant breeders. In this study, we staged anthers by development and performed a time-series analysis of sRNA and RNA transcripts expressed in anther development, in both wheat and barley. We describe a novel group of 24-nt phasiRNAs specifically expressed in pre-meiotic anthers.

## RESULTS

### A Developmental Time Series of Staged Wheat and Barley Anthers

Anthers develop from undifferentiated meristematic cells into an organized set of tissues with a plethora of functions. To identify pre-meiotic, meiotic and early post-meiotic stages of anther development in wheat and barley, anthers were dissected, fixed and processed for resin embedment and cross-sectioned. The developmental progression of meiosis was examined at 12 time points corresponding to 0.2 mm to 3.0 mm long anthers in bread wheat (cv. Fielder) and two barley varieties: Golden Promise (2-row; spring) and Morex (6-row; spring). Figure 1 shows a representative anther lobe for each sample and developmental stage. We observed a similar developmental progression between the two barley varieties as well as between wheat and barley. Cell fate specification was observed in 0.2 mm and 0.4 mm anthers, prior to observing (by 0.6 mm stage), the four distinct cell layers, namely the epidermis, endothecium, middle layer and tapetum (Figure 1). By 0.6 mm anthers, the microspore mother cells develop from sporogenous tissues. Subsequently, meiosis I and II occur in anthers, at the 0.8 mm to 1.4 mm stages. The tapetum was at its largest size (cell width) from 0.8 mm and 1.0 mm, which correlates to meiosis I, prior to degradation, until it was almost fully gone at the 1.6 mm to 1.8 mm stage, corresponding to the stage of vacuolated microspores. The vacuolated microspore develops into a mature pollen grains by 1.6 mm to 3.0 mm, when dehiscence occurs and pollen is released. Based on previous observations, we defined anthers to be pre-meiotic at ≤ 0.6 mm, meiotic from 0.8 mm to 1.4 mm, and post-meiotic by 1.6 mm. Thus, we sequenced RNA (sRNA and transcripts) of three pre-meiotic (0.2 mm, 0.4 mm and 0.6 mm), three meiotic (0.8 mm, 1.0 mm and 1.4 mm) and one post-meiotic (1.8 mm) anther stages.

**Figure 1.**
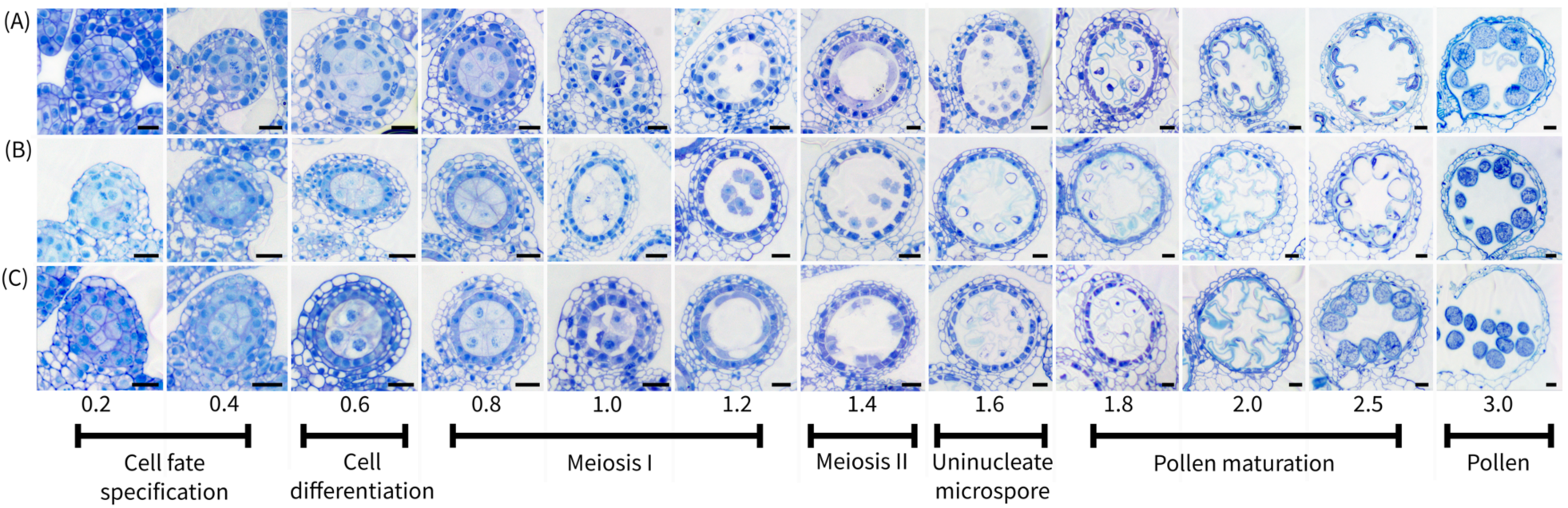
Transverse sections of wheat and barley anthers from 0.2 mm to the pollen stage. Anthers were obtained from the wheat cv. Fielder (A) and barley cv. Golden Promise (B) and Morex (C). Anthers were fixed with a 2% paraformaldehyde:glutaraldehyde solution and embedded using the Monostep HM20 polar resin, sectioning at 1.0 µm and stained using 1.0% toluidine blue O. Black scale bars correspond to 20 µm.

### A Comprehensive Annotation of Genomic Components Regulating PhasiRNA Biogenesis

To investigate the dynamics of phasiRNA biogenesis, regulation and accumulation in anthers, we deeply sequenced 126 libraries, evenly distributed across sRNA and poly(A) RNA from seven sequential stages of one bread wheat (cv. Fielder) and two barley (cv. Golden Promise and Morex) varieties. PhasiRNA biogenesis requires *PHAS* precursor transcripts, miRNA triggers and protein products of the *RDR, DRB, DCL* and *AGO* gene families. To provide a comprehensive overview of phasiRNA regulation in wheat and barley anthers, we performed an annotation of all these components and identified those expressed in anthers.

We first examined the miRNA triggers miR2118 and miR2275, as both families are required to cleave 21-*PHAS* and 24-*PHAS* precursor transcripts, respectively. Supplementary Table 1 details the coordinates and abundances of miRNAs annotated from anther tissues. A total of 240 miRNAs were annotated in wheat, of which 71 loci generating miR2118 and 15 loci of miR2275 were identified (Supplementary Table 1). In barley, 100 miRNAs were annotated of which 12 loci generating miR2118 and four loci of miR2275 were identified. A comparative analysis of miRNA triggers across the wheat sub-genomes revealed a bias in their distribution across homoeologous chromosomes. First, no miR2118 loci were observed on any homoeologs for chromosomes 3, 6 and 7 (Supplementary Figure 1a to 1c). Second, we observed that 31 of the 71 miR2118 loci (43.7%) came from the B sub-genome (Supplementary Table 1). That bias was mainly due to three clusters of miR2118 observed on chromosomes 1B, 4B and 5B, in which five, nine and nine miR2118 loci, respectively, were annotated, while only one cluster of four loci was observed on chromosome 5 of the A and D homoeologs (Supplementary Figure 1a to 1c). In barley, no miR2118 loci were detected on chromosomes 3H, 5H, 6H and 7H, although one cluster containing four miR2118 loci was observed on chromosome 4H (Supplementary Figure 1d). We did not observe any significant miR2275 clusters in wheat and barley, observing only 15 and four loci, respectively. Even so, the number of miR2275 loci differed between wheat homoeologous chromosomes; we annotated six, three, and six miR2275 loci on sub-genomes A, B and D, respectively (Supplementary Table 1; Supplementary Figure 1a to 1c). In contrast to miR2118, most of miR2275 loci were observed on wheat homoeologs of chromosomes 3 and 7 of sub-genomes A and D. Thus, our results show that the wheat sub-genomes unequally contribute miR2118 and miR2275 loci, as they are mainly located on the B or A and D sub-genomes, respectively. In brief, phasiRNA biogenesis can occur since the miRNA families triggering cleavage of *PHAS* transcripts were detected in anthers of both wheat and barley.

AGO proteins are required to load miRNA triggers and initiate *PHAS* transcript cleavage in addition to loading 21-nt or 24-nt phasiRNAs after their biogenesis. Although Komiya et al. (2014) identified the binding partner of 21-nt phasiRNAs, a binding partner for 24-nt phasiRNAs remains elusive. Thus, we carefully annotated all genes encoding AGO proteins in wheat and barley genomes and identified the *AGO* genes that were expressed in anthers. We found that wheat and barley genomes, respectively, encode a total of 67 and 22 *AGO* gene copies representing nine distinct *AGO* genes for each species (Supplementary Table 2; Figure 2). Of the total sets of *AGO* gene copies that we identified, 47 and 20 were expressed in wheat and barley anthers, respectively, and covered all distinct *AGO* genes. Our *de novo* transcript assembly has allowed us to describe, for the first time, two and eight new *AGO* gene copies in wheat and barley, respectively (Supplementary Table 2). Among the novel AGO proteins discovered in barley, we have annotated new copies for *AGO1* (three copies), *AGO2* (two copies), *AGO5* (two copies) and *AGO10* (one copy).

**Figure 2.**
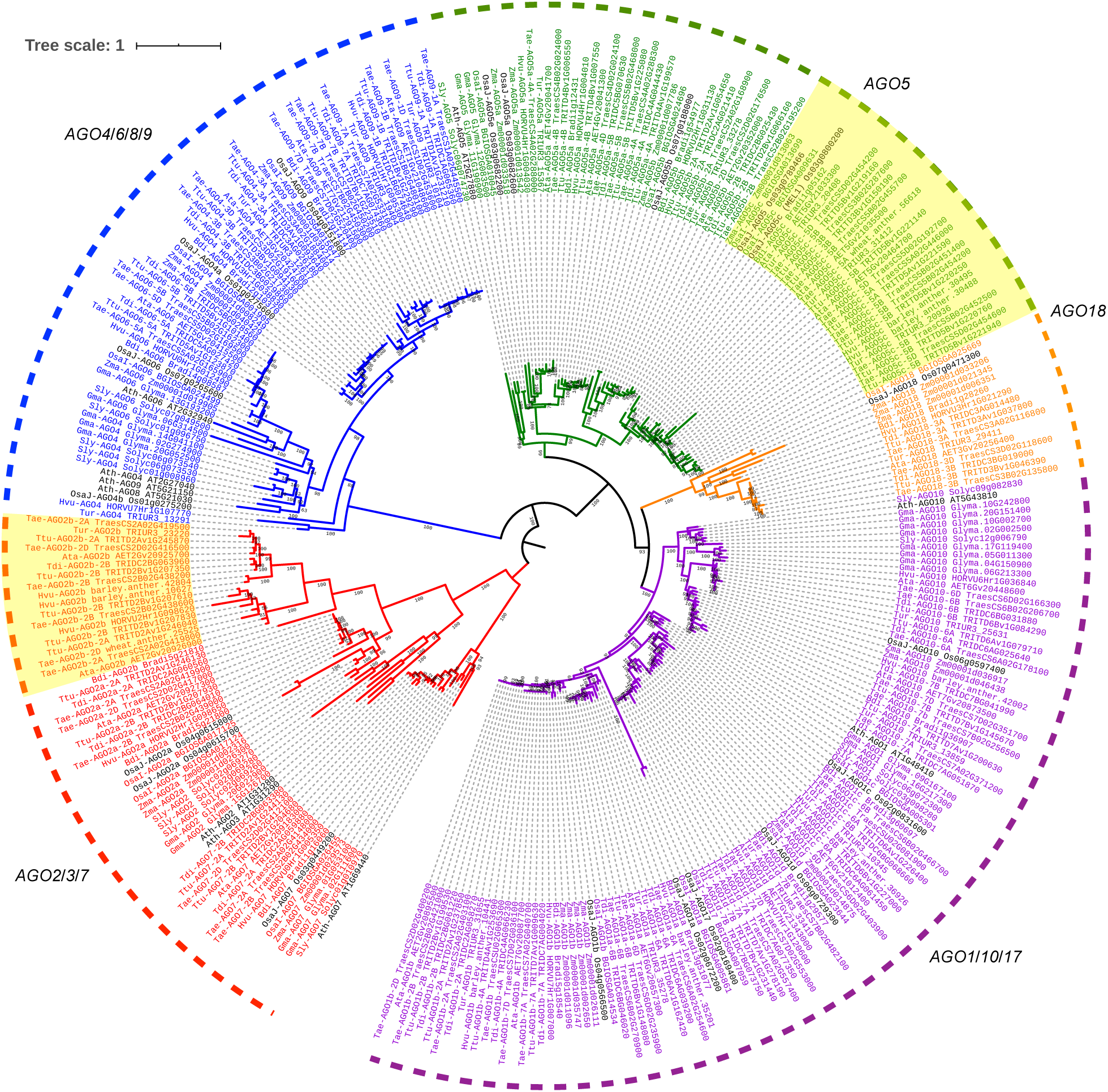
Phylogenic tree showing annotated wheat and barley orthologous genes to Argonaute (AGO) proteins. Clades of AGO protein were indicated. Yellow boxes highlight two groups of expended AGO proteins in wheat and barley. Protein orthologous groups were identified using OrthoFinder and SonicParanoid. Protein alignment and phylogeny analysis were done using MUSCLE and IQ-TREE.

Some interesting observations emerged from our phylogenetic analysis. First, the number and type of AGO proteins encoded by wheat and barley genomes were consistent if we take into account the hexaploid nature of wheat that results in a triplication of each *AGO* gene (Supplementary Table 2; Figure 2). Second, most of the *AGO* genes annotated in bread wheat were also identified in wheat-related ancestors carrying the A, B and D genomes. Third, the diversity of wheat, barley and other grasses is reflected in the evolutionary relationships of the *AGO1*/*2*/*5* genes (Figure 2). We observed a division of AGO1 protein in two sub-clades grouping AGO1a/b separate from AGO1c/d (Figure 2). Additionally, we observed an expansion of AGO1 proteins, duplicated from AGO1b, expanding the AGO1 clade to five copies in wheat, *T. dicoccoides* and barley, compared to four copies in the *O. sativa* or *B. distachyon* genomes. Compared to other grasses, an expansion of some *AGO2* and *AGO5* genes was also observed (Supplementary Table 2; Figure 2). In wheat, *AGO2b* was duplicated in each sub-genome expanding the number of copies of that *AGO* gene to six. In barley, we annotated three copies of the *AGO2b* gene. Compared to other grass species, the number of copies of *AGO5a*/*b*/*d* is the same. However, *AGO5c* was triplicated in wheat, bringing the total number of copies to 10, while in barley, one *AGO5c* duplication event was observed. Most *AGO* genes mentioned above were expressed in anthers of both species.

RDR, DRB and DCL are three other key proteins involved in phasiRNA biogenesis, thus, we annotated these families as well (Supplementary Table 2; Supplementary Figure 2a, b and c). The RDR6 protein synthesizes the second RNA strand of *PHAS* transcripts cleaved by AGO proteins. We found that the *RDR6* gene is duplicated in the barley genome. In contrast, we annotated only one *RDR6* gene in the wheat genome which is on chromosome 3 of the B and D homoeologs but not on the A homoeolog inherited from *T. urartu*. Curiously, we annotated, respectively, two and four *RDR6* genes in *T. urartu* and *T. dicoccoides* genomes, similar to barley, suggesting an incomplete annotation of *RDR* genes in the wheat genome. Unfortunately, we did not capture new transcripts for RDR6 protein in our *de novo* transcriptome assembly. Among other *RDR* genes, *RDR1* and *RDR3* are duplicated in both wheat and barley. In barley, all *RDR1* and *RDR3* copies were expressed in anthers. In wheat anthers, five of six *RDR1* copies were expressed in contrast to *RDR3*, for which only two of seven copies were expressed. Another group of essential proteins involved in phasiRNA biogenesis are the DCL and DRB proteins that, together, process a recursive slicing of 21- or 24-nt phased small RNA duplexes. A total of 15 and five gene copies, covering five distinct DCL proteins, were annotated in wheat and barley, respectively, which is exactly the same number as *DCL* genes in rice and maize (each having five copies). In wheat, each copy of these genes is evenly distributed across the A, B and D sub-genomes. Additionally, all annotated *DCL* genes were expressed in wheat and barley anthers including *DCL4* and *DCL5* genes that are known to be involved in, respectively, 21-nt and 24-nt phasiRNA biogenesis. A total of 25 and eight gene copies, covering five distinct DRB proteins, were annotated in wheat and barley, respectively. Both *DRB1* and *DRB2* were respectively duplicated and triplicated in both wheat and barley. DRB4 is a distinct DRB protein clade since most of its members carry three or four double-stranded RNA-binding motifs (DSRM) rather than two domains like other DRB proteins. Thus, DRB4 might have a distinct functional activity compared to other DRB proteins.

In summary, we characterized the set of phasiRNA biogenesis components in wheat and barley. We showed that all the machinery needed to process phasiRNAs is expressed in anthers of both species and, thus, phasiRNA biogenesis and functional activity should also be active.

### Myriad Stage-Specific phasiRNAs in Wheat and Barley Anthers

To allow accurate and sensitive identification of phasiRNAs that accumulate in anthers, we analyzed the 63 sRNA libraries generated from seven sequential stages of anther development. We identified a total of 12,821 and 2,897 *PHAS* loci in wheat and barley (detailed in Supplementary Table 3), respectively. Of the *PHAS* loci identified in barley, 2,699 and 2,824 were observed in cv. Golden Promise and Morex, respectively, of which 90.7% overlapped between the two varieties (Table 1). *PHAS* loci annotated in the wheat genome were fairly evenly distributed among homoeologous chromosomes; the A, B and D wheat sub-genomes, respectively, encompassed 4,209, 4,079 and 3,930 loci (Table 1). We detected 2.4-fold more 21-*PHAS* than 24-*PHAS* loci in these genomes. That ratio was consistent between wheat and barley genomes, or barley cultivars, or wheat sub-genomes (Table 1). *PHAS* loci were fairly evenly distributed to all chromosomes over loci corresponding to euchromatic genomic regions in both the wheat and barley genomes (Supplementary Figure 1). Overall, fewer than 20 sRNA loci overlapped repetitive regions across nearly 3,000 loci in barley, and not one was detected that overlapped repetitive regions in wheat across almost ∼13,000 loci. This characteristic distinguishes the group of 24-nt phasiRNAs from the plant DCL3-dependent siRNAs, the 24-nt heterochromatic siRNAs, which are mostly derived from repetitive elements, primarily transposable elements.

**Table 1.** Total number of 21- and 24-*PHAS* loci annotated in the wheat and barley genomes.

| *PHAS* loci | **Wheat** |  |  |  |  | **Barley** |  |  |
| --- | --- | --- | --- | --- | --- | --- | --- | --- |
|  | Total | AA* | BB* | DD* | Unordered** | Total | Golden Promise | Morex |
| Total number of loci | 12,821 | 4,209 | 4,079 | 3,930 | 603 | 2,897 | 2,699 | 2,824 |
| 21-PHAS | 9,073 | 2,994 | 2,852 | 2,786 | 441 | 2,022 | 1,902 | 1,984 |
| 24-PHAS | 3,748 | 1,215 | 1,227 | 1,144 | 162 | 875 | 797 | 840 |
| Ratio | 2.4 | 2.5 | 2.3 | 2.4 | 2.7 | 2.3 | 2.4 | 2.4 |

\* The sub-genomes in which *PHAS* loci were identified.
\*\* *PHAS* loci located on unordered DNA scaffolds of the wheat genome. The sub-genome of origin could not be designated.

We observed a distinct temporal accumulation of 21-nt and 24-nt phasiRNAs in anther development. In Figure 3, we show that 21-nt phasiRNAs accumulate in barley anthers by the 0.2 mm to 0.6 mm stages, during cell fate specification (in 0.2 mm and 0.4 mm) and the differentiation of the four cell layers (0.6 mm), prior to decreasing in the early meiotic stage (0.8 mm or 1.0 mm). At their peak in quantity and diversity (in 0.2 mm to 0.8 mm), 21-nt phasiRNAs represented more than 90% of all 21-nt sRNAs detected in anthers; this was much higher than the 60% peak proportion of 21-nt reproductive phasiRNAs observed in maize (Zhai et al., 2015). We observed a different phasiRNA accumulation pattern for 24-nt phasiRNAs; in this case, hundreds of 24-nt phasiRNAs remained undetectable until the 0.8 mm stage, when anthers enter early meiotic stage (Figure 3). At their peak, corresponding to 0.8 mm to 1.4 mm anthers (with a maximum at 1.0 mm), 24-nt phasiRNAs reached 93% of all 24-nt sRNAs detected in anthers. This was again substantially greater than the 64% peak proportion observed in maize (Zhai et al., 2015). Although their abundance stays detectable, in barley, we observed a significant decrease in 21-nt and 24-nt phasiRNAs abundance in post-meiotic anthers (1.8 mm stage), as well as a decrease in their proportion of all 21-nt or 24-nt sRNAs, respectively. After their peak of accumulation, the abundance of wheat 21-nt and 24-nt phasiRNAs decreased less than observed in barley.

**Figure 3.**
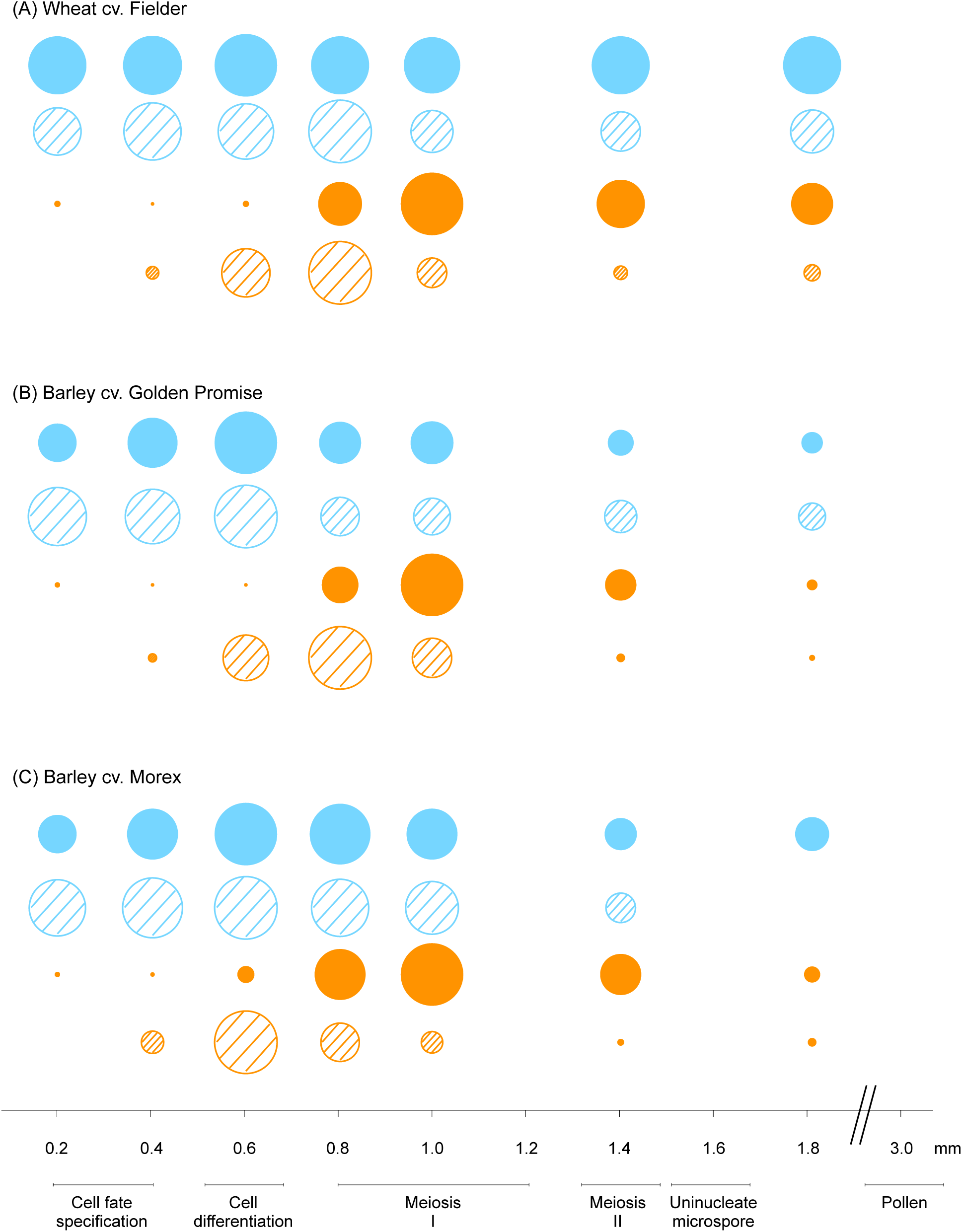
The relative accumulation of reproductive phasiRNAs and their miRNA triggers. The triggers of 21-nt (blue circle) and 24-nt (orange circle) phasiRNAs are respectively miR2118 (blue hashed-circle) and miR2275 (orange hashed-circle). These were characterized during the development of anthers in wheat cv. Fielder (A) and barley cv. (B) Golden Promise and (C) Morex.

PhasiRNA biogenesis requires miRNA triggers of both the miR2118 and miR2275 families to cleave 21-*PHAS* and 24-*PHAS* precursor transcripts, respectively. The abundance of miR2118 family members peaked in pre-meiotic stages (0.2 mm to 0.6 mm) and then decreased in early meiotic (0.8 mm or 1.0 mm) stages (Figure 3). Thus, good synchrony was observed between abundance peaks of 21-nt phasiRNAs and miR2118. In contrast, expression of miR2275 family members peaked at 0.6 mm or 0.8 mm, prior to a drastic decrease that occurred before the burst of 24-nt phasiRNAs that reached their peak at 1.0 mm (Figure 3). Collectively, miRNA triggers and phasiRNA products showed a coordinated accumulation pattern in wheat and barley varieties suggesting there is good conservation in the regulation of 21-nt and 24-nt phasiRNA pathways during anther development in the *Triticeae* tribe.

### Pre- and Post-Meiotic Accumulation of 24-nt PhasiRNAs

Although the large majority of miR2275 and 24-nt phasiRNAs reach their peak in anthers at 0.8 mm to 1.0 mm, we examined the abundance of all 24-*PHAS* loci. We discovered three distinct accumulation patterns of 24-nt phasiRNAs in wheat and barley. For example, in wheat, the group (group “a”) that accumulates as previously described, 1,611 24-*PHAS* loci (42.9% of total 24-*PHAS* loci) were supported by 30.5 million (M) reads which represent ∼81% of all 24-nt phasiRNA reads (Figure 4a). The 24-nt phasiRNAs of this cluster are almost completely absent (median ≤ 5 transcript per millions of mapped reads (TPMM)) in barley cv. Morex and wheat anthers smaller than 0.6 mm, and in barley cv. Golden Promise anthers smaller than 0.8 (Figure 4b). At their peak of abundance (by 1.0 mm anthers), the median abundance increased greater than 1000-fold, when anthers reached the meiotic stage. Among putative *PHAS* precursor transcripts of this group, we found one conserved motif that matched perfectly with miR2275 and was present in, as an example in wheat, 84.6% of all *PHAS* precursors (Figure 4c).

**Figure 4.**
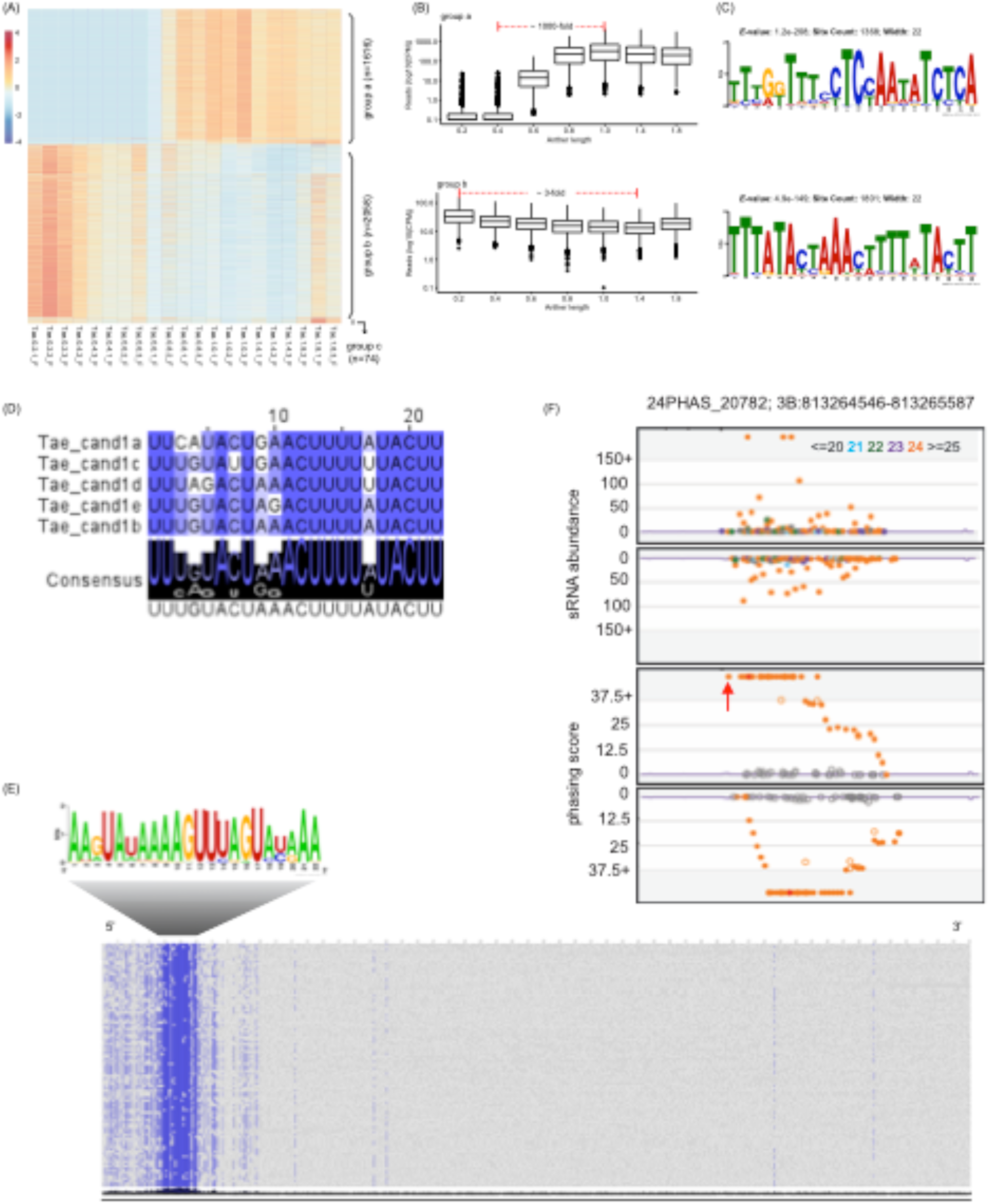
Pre-meiotic 24-nt reproductive phasiRNAs found in the wheat cv. Fielder. We show the accumulation pattern of 24-nt phasiRNAs illustrated by heatmap (A) and box plot (B) graphs. (C) The two conserved motifs found in putative *PHAS* transcripts for pre-meiotic and meiotic phasiRNAs. (D) Alignment of candidate sRNA triggers for pre-meiotic 24-*PHAS* loci. The degree of conservation is denoted by density of blue color and consensus sequence is shown with sequence logos. (E) Sequence logo denoting conservation of target site (Above). Nucleotide sequence alignment for 100 pre-meiotic 24-*PHAS* loci showing the conservation of the candidate miRNA triggers target sites (Below). (F) Two tracks showing the abundance (RP10M) of small RNAs in both strands of a representative pre-meiotic 24-*PHAS* loci (Above). sRNA sizes are indicated by different color as shown. Two tracks showing the phasing score of each read for both strands in this locus (Below). The red arrow indicates the candidate cleavage site from where the first phasiRNA is generated.

The accumulation pattern for the second group (group “b”) of 24-*PHAS* loci is quite distinctive from group “a”. We observed a peak of abundance at the earliest stage of anther development (0.2 mm) with abundances gradually decreased through development for both barley varieties and in wheat. In wheat, this group consists of 2,058 24-*PHAS* loci (54.9% of total 24-*PHAS* loci) and is supported by ∼19% of all 24-nt phasiRNA reads (7.4 M reads). Thus, their abundance peaks in pre-meiotic anthers and decreases ∼3-fold by meiotic stages. Among putative *PHAS* precursor transcripts in this group, using wheat as an example, we found one conserved motif with no homology to miR2275, or any other known miRNA, which was present in 87.5% of all *PHAS* precursors in group “b”.

To identify the sRNA capable to trigger the 24-nt pre-meiotic phasiRNAs, we searched for sRNA having a strong homology to the conserved motif within our sequenced sRNA libraries. We could identify five, and four, candidate sRNA triggers in wheat and barley, respectively (Figure 4d). These candidate triggers are 22-nt in length and start with a 5’ uridine but we could not classify them as miRNA since no hairpin structure could confirmed. The lack of a canonically-folded putative precursors of these sRNA triggers may be due to an incomplete genome sequence in the region at which these sRNAs are encoded. However, the candidate triggers aligned exactly to the position of the conserved motif found in 24-nt pre-meiotic phasiRNAs precursors (Figure 4e). Using wheat as an example, a target prediction analysis predicted 1,524 (74.1% of all) sRNA-target interaction modules. At a target score cut off of 4, candidate sRNA triggers were appropriately matched to direct cleavage of 790 (38.4%) 24-nt pre-meiotic *PHAS* transcripts. The read distribution and phasing score of a representative 24-nt pre-meiotic phasiRNA is displayed in Figure 4f. Similar findings were observed in both varieties of barley.

We observed a third group (group “c”) in the two barley varieties, but this group was less obvious in wheat. Only 19 and 20 *PHAS* group “c” loci were detected for barley cv. Golden Promise and cv. Morex, respectively. Although these loci were detectable in pre-meiotic and meiotic anthers, their peak in abundance was in the post-meiotic stage of anthers, by 1.8 mm (Figure 4). Even though few *PHAS* loci displayed this pattern of read accumulation, representing a low proportion of the total 24-nt phasiRNAs, this cluster accumulating in post-meiotic stages may have a biological function in gametogenesis. In brief, our studies have shown that there are two distinct types of 24-phasiRNAs that peak in pre-meiotic and meiotic wheat and barley anthers while a third category of phasiRNA accumulates in post-meiotic barley anthers.

### Stage-Specific PhasiRNAs are Co-expressed with Specific DRB, DCL and AGO Genes

The distinct temporal accumulation of 21-nt and 24-nt phasiRNAs requires precise regulation of *PHAS* precursor transcription and of the biogenesis components of phasiRNA pathways. Transcripts assembled from 63 poly(A) RNA libraries were used to perform a co-expression analysis to identify the genes that might regulate the phasiRNA groups described above. First, 12, 14, and 15 gene modules were identified in wheat and barley varieties Golden Promise and Morex, respectively (Supplementary Figure 3). Relative to meiotic progression, some modules were associated with distinct phases of anther development. In wheat, for example, a total of two (modules 3 and 5), three (modules 1, 2 and 7), and three (modules 4, 6 and 8) expression modules were strictly associated with pre-meiotic, meiotic and post-meiotic anthers, respectively (Supplementary Figure 3). Thus, we analyzed genes of these modules correlated with phasiRNA accumulation, with an emphasis on *DCL* and *DRB* genes, and for those encoding candidate AGO binding partners of phasiRNAs.

These genes are summarized in Table 2 (as well as in Supplementary Figure 3), classified into pre-meiotic, meiotic or post-meiotic stages. Most of these genes show the same co-expression pattern, relative to meiotic progression, for the wheat and the two barley varieties (Table 2; Supplementary Figure 3). Thus, we focused on those showing shared expression patterns. In pre-meiotic anthers, many *AGO* genes, namely, *AGO1b*/*d, AGO5c, AGO7, AGO9*, and *AGO10*, in addition to four *DCL* (*DCL1, DCL2, DCL3* and *DCL4*) and three (*DRB1, DRB2c* and *DRB5*) genes, were co-expressed (Table 2; Supplementary Figure 3). These genes were co-expressed with two groups of phasiRNAs, namely, 21-nt phasiRNAs and pre-meiotic 24-nt phasiRNAs. Meiotic 24-nt phasiRNA expression correlated to numerous AGO genes, namely, *AGO2b, AGO5b, AGO6, AGO18* as well as to *DCL5* and *DRB4* genes (Table 2; Supplementary Figure 3). We found a strong correlation between genes mentioned above and anther developmental stages; due to the hexaploid nature of wheat, multiple copies of the same gene showed the same pattern. For example, the wheat genome encodes a total of 10 AGO5c proteins for which seven genes were expressed in anther and all the corresponding genes were co-expressed with the pre-meiotic stage of anthers. In wheat, all homoeologous copies of the genes reported above showed the same expression pattern in pre-meiotic (*AGO1b*/*d, AGO5c, AGO9, AGO10, DCL4*) and meiotic (*AGO2b, AGO5b, AGO6, AGO18, DCL5, DRB4*) anthers regardless of their total number of copies (three, six, or nine). In barley, the same coherence was observed. For instance, the *AGO2b* gene in barley was triplicated in its genome and all copies were co-expressed in meiotic anthers. In summary, we found that the accumulation of stage-specific reproductive phasiRNAs correlates to specific and distinct *DCL, DRB* and *AGO* genes.

**Table 2.** Co-expressed genes correlated to anther developmental phases for *DCL, DRB* and *AGO* genes.

|  | ***Pre-meiotic*** | ***Meiotic*** | ***Post-meiotic*** |
| --- | --- | --- | --- |
| Wheat<br>(Fielder) | Tae-AGO1b/d<br>Tae-AGO5c<br>Tae-AGO9<br>Tae-AGO10<br>Tae-DCL1<br>Tae-DCL2<br>Tae-DCL4<br>Tae-DRB1<br>Tae-DRB2a/c<br>Tae-DRB5 | Tae-AGO2b<br>Tae-AGO5b<br>Tae-AGO6<br>Tae-AGO18<br>Tae-DCL5<br>Tae-DRB4 | Tae-AGO1c |
| Barley<br>(Golden Promise) | Hvu-AGO1b/c/d<br>Hvu-AGO4<br>Hvu-AGO5c<br>Hvu-AGO7<br>Hvu-AGO9<br>Hvu-AGO10<br>Hvu-DCL1<br>Hvu-DCL2<br>Hvu-DCL3<br>Hvu-DCL4<br>Hvu-DRB1<br>Hvu-DRB2a/c<br>Hvu-DRB5 | Hvu-AGO2b<br>Hvu-AGO5b<br>Hvu-AGO6<br>Hvu-AGO18<br>Hvu-DCL5<br>Hvu-DRB4 | Hvu-AGO2a<br>Hvu-AGO5a<br>Hvu-DCL4 |
| Barley<br>(Morex) | Hvu-AGO1b/d<br>Hvu-AGO4<br>Hvu-AGO5c<br>Hvu-AGO7<br>Hvu-AGO9<br>Hvu-AGO10<br>Hvu-DCL2<br>Hvu-DCL3<br>Hvu-DCL4<br>Hvu-DRB1<br>Hvu-DRB2c<br>Hvu-DRB5 | Hvu-AGO2b<br>Hvu-AGO5b<br>Hvu-AGO6<br>Hvu-AGO18<br>Hvu-DCL1<br>Hvu-DCL5<br>Hvu-DRB4 | Hvu-AGO1c<br>Hvu-AGO2a<br>Hvu-DRB2b<br>Hvu-DCL4 |

Note: Gene IDs can be found in Supplementary Table 2

## DISCUSSION

A detailed understanding of morphological development of anthers is necessary to enable studies of the molecular pathways governing their development. Although the morphological development of wheat (Browne et al., 2018; Shunmugam et al., 2018) and barley (Waddington et al., 1983; Gómez and Wilson, 2012) anthers was described previously, in this study we staged anthers of commonly studied varieties, as they differ from those previously characterized. Additionally, we increased the resolution of developmental stages to every 0.2 mm of development, which provided greater detail and augmented previous results in wheat (Browne et al., 2018) and barley (Gómez and Wilson, 2012). Relative to their length, an approach that aids harvesting of anthers for future molecular studies, we identified pre-meiotic, meiotic and post-meiotic anthers. We detected the initiation of meiosis I by 0.8 mm anthers, and meiosis II in 1.4 mm anthers in wheat cv. Fielder and both barley varieties. In wheat, our observations were consistent with Shunmugam et al. (2018), who described each phase of meiosis I in wheat anthers of the Chinese Spring cultivar, between 0.7 mm to 1.2 mm. Together, our results, and those reported by Shunmugam et al. (2018), suggest a similar developmental kinetic of anthers between, or within, wheat and barley species. However, additional staging of more varieties of each species may allow more precise correlation of meiosis progression with anther length. To enable this, we performed microscopy analyses to identify pre-meiotic (0.2 mm, 0.4 mm and 0.6 mm), meiotic (0.8 mm, 1.0 mm and 1.4 mm) and post-meiotic (1.8 mm) anthers for which we then investigated accumulation patterns of RNAs, including reproductive phasiRNAs.

In recent years, two classes of pre-meiotic (21-nt) and meiotic (24-nt) phasiRNAs and their patterns of accumulation in anthers, have been described in maize (Zhai et al., 2015) and rice (Johnson et al., 2009; Fei and al., 2016). Their precise function remains unclear but some studies have shown that they play a critical role in supporting male fertility (Fan et al., 2016; Teng et al., 2020). Their important role in anthers underpins our current study to characterize phasiRNAs in wheat and barley anthers, for which there is, to date, only one published sRNA analysis in wheat but nothing in barley. In this study we annotated a total 12,821 *PHAS* loci in the wheat genomes and observed a ratio of 2.4-fold 21-*PHAS* to 24-*PHAS* loci. Earlier this year, Zhang et al. (2020) revisited public sRNA data generated from reproductive and non-reproductive wheat tissues. Their goal was to annotate *PHAS* loci in the wheat genome and compare the *PHAS* distribution among wheat sub-genomes. In non-reproductive wheat tissues, they reported few phasiRNAs (limited 21-nt phasiRNAs and no 24-nt phasiRNAs), illustrating the reproductive nature of phasiRNAs. At the same threshold that we used, Zhang et al. (2020) annotated a total of 8,450 non-redundant *PHAS* loci and a ratio of 1.3-fold 21-*PHAS* to 24-*PHAS* loci. Overall, we discovered 4,371 more *PHAS* loci, perhaps due to our more extensive sampling. Most of the *PHAS* loci that Zhang et al. (2020) did not annotate were 21-nt phasiRNAs which mainly accumulate in pre-meiotic anthers. We attribute our detection of those phasiRNAs to our analysis of isolated anthers dissected from the spike, whereas Zhang et al. (2020) examined sRNAs in spikes rather than anthers, limiting their ability to annotate these several thousands of *PHAS* loci.

We also annotated 2,897 *PHAS* loci in the barley genome. Due to the hexaploid nature of the wheat genome, we might expect to find 3-fold more *PHAS* loci in wheat than in barley. However, we observed ∼4.4-fold more *PHAS* loci in wheat than in barley. When considering the larger depth of sequencing performed for wheat libraries compared to barley, respectively, 2.7-fold (for barley cv Morex) or 3.4-fold (for barley cv. Golden Promise), it is possible or even probable that we missed perhaps a few hundred *PHAS* loci in the barley genome. Regardless, the number of *PHAS* loci we annotated in the wheat and barley genomes is much higher in comparison to the 639 loci previously annotated in the maize genome (Zhai et al., 2015). We detected 2.4-fold more 21-*PHAS* than 24-*PHAS* loci in wheat and barley cultivars, a ratio similar to maize, for which Zhai et al. (2015) reported a ratio of 2.6-fold 21-*PHAS* to 24-*PHAS* loci. Fei et al. (2016) annotated 1,843 21-*PHAS* loci in the rice genome, a similar number to the 2,022 21-*PHAS* loci that we annotated in the barley genome in this study. Yet, Fei et al. (2016) annotated just 50 24-*PHAS* loci, which is extremely low compared to the number that we annotated in the wheat (3,748) and barley (875) genomes. Fei et al. (2016) analyzed 24-nt phasiRNA in 0.40 to 0.45 mm rice anthers, corresponding to an early meiotic stage (Zhang et al., 2011). Thus, it is possible that the lower number of 24-*PHAS* loci reported previously in the rice genome, may be due to sampling only at earlier, pre-meiotic, stages of development (Fei et al., 2016). When comparing the total number of *PHAS* loci in genome of maize, rice, barley and wheat, we notice an expansion of reproductive *PHAS* loci in genomes of *Poaceae* subfamilies from *Panicoideae* to *Oryzoideae* and to *Poideae*.

In addition to the two classes of pre-meiotic (21-nt) and meiotic (24-nt) phasiRNAs, previously described in maize and rice anthers (Zhai et al., 2015; Fei et al., 2016), we also described a new group of 24-nt phasiRNAs that accumulate in pre-meiotic anthers, more like the known 21-nt phasiRNAs. We observed these novel pre-meiotic 24-nt phasiRNAs in both wheat and barley anthers, in which they represented more than 50% of all 24-*PHAS* loci that we annotated. Pre-meiotic 24-nt phasiRNAs have not been reported in either maize or rice, despite extensive sampling (i.e. Zhai et al., 2015). We re-analyzed these older sRNA data from maize anthers (Zhai et al., 2015) and we detected no evidence of pre-meiotic 24-nt phasiRNAs. This apparent absence of pre-meiotic 24-nt phasiRNAs in maize and rice suggests a divergence in grass species of the *Poideae* subfamily. Future studies on more grass subfamilies could shed light on the emergence and evolution of this new class of pre-meiotic 24-nt phasiRNAs.

Initiation of 24-nt meiotic phasiRNA biogenesis canonically occurs at a miR2275 target site (Johnson et al., 2009; Song et al., 2012a). We observed this conserved motif, homologous to miR2275, in *PHAS* precursors of meiotic 24-nt phasiRNAs. For the novel, pre-meiotic 24-nt phasiRNAs, we found that miR2275 is not expressed coincident with their accumulation in anthers, and no motif matching miR2275 was present in their *PHAS* precursors. We did identify, in these *PHAS* precursors, a well-conserved motif that lacked confident homology to known miRNAs registered on miRBase and to novel miRNAs annotated in wheat or barley. However, we identified a candidate sRNA trigger in wheat (five copies in the genome) and barley (four copies in the genome) with evidence supporting its role in triggering the cleavage of pre-meiotic 24-nt *PHAS* transcripts. The poor assembly at the genomic origin of these sRNAs does not allow us to annotate them as miRNAs, since we could not confirm a hairpin precursor sequence at these loci. Thus, it remains unclear how the biogenesis of these novel, pre-meiotic 24-nt phasiRNA is initiated. Rather than initiation via a miRNA-directed, AGO-catalyzed cleavage of a single-stranded *PHAS* precursor, phasiRNA biogenesis may be initiated by an as-yet unknown mechanism.

PhasiRNA biogenesis and regulation requires a set of specific proteins. Over the last decade, both the 21- and 24-nt phasiRNA pathways have been described. Functional studies have shown that generation of pre-meiotic 21-nt phasiRNAs is dependent on miR2118, RDR6, DCL4 and MEL1, and presumably a copy of AGO1 (Komiya et al., 2014; Nonomura et al., 2007; Song et al., 2012a; Song et al., 2012b). Fei et al. (2016) performed a co-expression analysis in pre-meiotic rice anthers and reported that the expression pattern of *OsAGO1d* was the same as *OsAGO5c*, suggesting a functional connection between AGO1d and pre-meiotic phasiRNAs. Similarly, in this study, we observed the same expression pattern for the *AGO1b* and *AGO5c* genes in wheat and barley anthers. Furthermore, the wheat genome encodes a total of 10 *AGO5c* proteins of which seven are expressed in anthers. Concordantly, we found that all the gene copies showed the same expression in phase with the peak of 21-nt, pre-meiotic phasiRNA expression. Moreover, our analysis revealed that *AGO7* (only in barley anthers), *AGO9* and *AGO10* were also co-expressed with *AGO1d* and *AGO5c*, although it remains unclear how these AGO proteins may be involved. Perhaps their role is in a developmental process such as the regulation of shoot apical meristems, as by *AGO10* (Zhang et al., 2015), or other functions in anther development, independent of a role in the reproductive 21-nt phasiRNA pathway.

Numerous studied have described the 24-nt phasiRNA pathway as dependent on miR2275, *RDR6* and *DCL5*, presumably as well as a copy of *AGO1* (Song et al., 2012a; Song et al., 2012b; Zhai et al., 2015; Fei et al., 2016; Teng et al., 2020). Until now, no functional studies have identified an AGO protein as the binding partner of 24-nt phasiRNAs. However, some promising candidates *AGO* genes have been proposed in maize and rice. In rice, Fei et al. (2016) showed that the expression of *AGO2b, AGO5b* and *AGO18* correlated with the peak of 24-nt meiotic phasiRNA accumulation. We made the same observation in wheat and barley anthers. We also found a strong correlation of *AGO6* expression with the burst of 24-nt phasiRNA in meiotic anthers. Additionally, the normalized abundance of *AGO6* expression is ∼1.5-fold higher than *AGO5b* or *AGO18*, and 5 to 10-fold higher than *AGO2b*. Similar to *AGO4* and *AGO9*, the AGO6 protein was described as a binding partner of 24-nt heterochromatic siRNAs (Borges and Martienssen, 2015) associated with RNA-directed DNA methylation (RdDM). AGO6 thus represents a promising candidate as a binding partner to 24-nt meiotic phasiRNAs.

Among the *AGO* genes co-expressed with pre-meiotic 24-nt phasiRNAs, *AGO9* is of particular interest, because in Arabidopsis female reproductive development, *AGO9* is a binding partner to heterochromatic 24-nt sRNAs in the silencing of transposable elements (TEs) (Olmedo-Monfil et al., 2010; Zhang et al., 2015). In both Arabidopsis and maize, *AGO9* represses germ cell fate in the somatic cells surrounding the precursors of the gametic cells (Olmedo-Monfil et al., 2010; Singh et al., 2011; Zhang et al., 2015). A loss-of-function *ago9* mutant in maize exhibits unreduced (diploid) gametes, abnormal chromatin condensation during meiosis, and hypomethylated repeats in centromeres (Singh et al., 2011; Zhang et al., 2015). Together, the binding affinity of *AGO9* with 24-nt heterochromatic sRNAs, its role in the female reproductive development plus the co-expression of *AGO9* with pre-meiotic 24-nt phasiRNAs (in wheat and barley) highlight *AGO9* as a strong candidate for the binding partner of pre-meiotic 24-nt phasiRNAs.

## CONCLUSION AND PERSPECTIVES

An efficient and cost-effective hybrid seed program has been difficult to develop in wheat and barley since they are self-fertilized species. The development of genic male-sterile lines may serve as the basis for the development of such a breeding program, via control of the production of pollen. In rice, such breeding programs may utilize photoperiod- or temperature-sensitive genic male-sterile lines (Ma and Yuan, 2015). Perturbed individual 21-nt reproductive phasiRNAs underlies this trait in rice (Ding et al., 2012; Fan et al., 2016). In maize, loss of the 24-nt reproductive phasiRNAs also confers a conditional male sterile phenotype (Teng et al., 2020). To develop these types of genic male-sterile lines in wheat and/or barley, it will be important to fully characterize the loci and their expression patterns during anther development that yield these reproductive phasiRNAs, and it will be important to understand the genes involved in their biogenesis.

We describe a catalog of *PHAS* loci in wheat and barley, at high resolution, over the course of anther development. We discovered a large cohort of pre-meiotic 24-nt phasiRNAs not previously observed in rice or maize. Questions remain, for example, which AGO proteins are the binding partners of pre-meiotic and meiotic 24-nt phasiRNAs. In the near future, more functional experiments are needed to fully characterize the 24-nt phasiRNA pathway and to precisely describe the biological function of both 21- and 24-nt reproductive phasiRNAs. Ideally, this additional knowledge will enable the modulation of male fertility with varied environmental conditions such as light or temperature.

## MATERIAL AND METHODS

### Plant Growth Conditions and Tissue Harvesting

Plants of bread wheat (*Triticum aestivum ssp. aestivum*) cv. Fielder (spring) and barley (*Hordeum vulgare ssp. vulgare*) cv. Golden Promise (2-row; spring) and Morex (6-row; spring) were grown in a greenhouse under conditions of 20/18°C day/night, 16/8h light/dark, and 50% relative humidity. For both species, we used a stereomicroscope (Mantis Elite-Cam HD, Vision Engineering, New Milford, CT, USA) to dissect anthers at the following lengths: 0.2 mm, 0.4 mm, 0.6 mm, 0.8 mm, 1.0 mm, 1.2 mm, 1.4 mm, 1.6 mm, 1.8 mm, 2.0 mm, 2.5 mm and 3.0 mm. Anthers were collected only from middle spikelets of the first three tillers. Only anthers from the first florets, in wheat, or the main spikelets, in barley, were dissected. For histological analysis, 10 anthers were collected per sample. For RNA analysis, per sample, a total of 125 (anther length <0.8 mm), 75 (anther length between 0.8 mm to 1.2 mm) and 50 (anther length >1.2 mm) anthers were collected for RNA isolation. Anthers were harvested in three replicates and, after dissection, samples were immediately frozen in liquid nitrogen and kept at −80°C prior to RNA isolation.

### Tissue Embedding and Microscopy

Fresh young spikelets were fixed (2% paraformaldehyde and 2% glutaraldehyde plus 0.1% Tween20 in 0.1M PIPES buffer at pH 7.4) overnight and dehydrated through a standard ethanol series (30%, 50%, 70%, 80%, 90%, 100% of cold EtOH) prior to being resin infiltrated and embedded using the Lowicryl Monostep Embedding Media HM20 (#14345, Electron Microscopy Sciences, Hatfield, PA, USA) using either heat or UV polymerization. Embedded tissues were sectioned at 1 µm using the Leica Ultracut UCT (Leica Microsystems Inc., Buffalo Grove, IL, USA) and stained using a 1.0% Toluidine Blue O dye (#26074-15, Electron Microscopy Sciences, Hatfield, PA, USA). Microscopy images were captured using a ZEISS Axio Zoom.V15 microscope using the PlanNeoFluar Z 2.3x/0.57 FWD 10.6mm objective lens with a magnification of 260X. Digital images were captured at 2584 × 1936 pixel resolution at 12 bit/channel.

### RNA Isolation, Library Construction and Sequencing

Total RNA was extracted with TRI Reagent (Sigma-Aldrich, St Louis, MO, USA) following the manufacturer’s instructions. RNA quality was assessed using an Agilent RNA 6000 Nano Kit with the Bioanalyzer 2100 (Agilent Technologies, Santa Clara, CA, USA). Only samples with an RNA integrity number ≥ 7.0 were kept for library construction. Small RNA libraries were prepared using the RealSeq-AC miRNA Library Kit for Illumina sequencing (Somagenics, Santa Cruz, CA, USA), following the manufacturer’s recommendations. Small RNA libraries were size-selected with the end product of ∼150 nt using a polyacrylamide/urea gel or the SPRIselect Reagent (Beckman Coulter Life Sciences, Indianapolis, IN, USA) magnetic beads for, respectively, wheat and barley. RNA-seq libraries were constructed using a NEBNext Ultra II Directional RNA Library Prep Kit for Illumina (#E7765L, New England Biosystems, Ipswich, MA, USA), following the manufacturer’s recommendations to prepare cDNA libraries having an insert of ∼200 bp or 300-350 bp for wheat and barley, respectively. Single-end sequencing was performed; 76 (3 lanes) and 151 cycles (3 lanes) for, respectively, sRNA-seq and RNA-seq libraries. The sequencing was generated on an Illumina NextSeq 550 instrument (Illumina Inc., San Diego, CA, USA), at the University of Delaware DNA Sequencing & Genotyping Center (Newark, DE, USA).

### Bioinformatics Analysis of sRNA-Seq Data

Using cutadapt v1.9.1 (Martin, 2011), sRNA-seq reads were pre-processed to remove the 3’ adapter; trimmed reads shorter than 15 nt were discarded. These cleaned reads were mapped to the wheat (Triticum_aestivum.IWGSC.dna.toplevel) and barley (Hordeum_vulgare.IBSC_v2.dna.toplevel) reference genomes (ftp://ftp.ensemblgenomes.org/pub/plants/release-44) using ShortStack v3.8.5 (Johnson et al., 2016) with following parameters: -mismatches 0, -bowtie_m 50, -mmap u, -dicermin 19, -dicermax 25, -foldsize 800 and -mincov 0.5 transcripts per million mapped (TPMM). Results reported by ShortStack were filtered to keep only clusters having a predominant RNA size observed between 20 and 24 nucleotides inclusively. We then annotated classes of miRNAs and phasiRNAs.

First, sRNA reads representative to each cluster were aligned to monocot-related miRNAs listed in miRBase release 22 (Kozomara and Griffiths-Jones, 2014; Kozomara et al., 2019) using ncbi-blastn v2.9.0+ (Camacho et al., 2009) with following parameters: -strand both, -task blastn-short, -perc_identity 75, -no_greedy and -ungapped. Homology hits were filtered and sRNA reads were classed as known miRNAs, based on the following criteria: (i) no more than four mismatches and (ii) no more than 2-nt extension or reduction at the 5’ end or 3’ end. Known miRNAs were summarized by family and when genomes contained multiple miRNA loci per family, miRNAs were ordered per chromosomal position and renamed base on these genomic positions. Small RNA reads with no homology to known miRNA were annotated for new miRNA using the *de novo* miRNA annotation performed by ShortStack. The secondary structure of new miRNA precursor sequences was drawn using the RNAfold v2.1.9 program (Lorenz et al., 2011). Candidate novel miRNAs were manually inspected and only those meeting criteria for plant miRNA annotations (Axtell and Meyers, 2018) were kept for downstream analyses. Then, remaining sRNA clusters were analyzed to identify phasiRNA-generating loci based on the ShortStack analysis report. sRNA clusters having a “Phase Score” exceeding or equal to 30 were considered as true positive phasiRNA loci. Genomic regions at these phasiRNAs were identified as *PHAS* loci and grouped in categories of 21- and 24-*PHAS* loci based on the length of phasiRNA derived from these loci.

### Identification of Candidate 24-nt Pre-Meiotic PhasiRNA Triggers

We used MEME v5.1.0 (Bailey et al., 2009; Bailey et al., 2015) to identify and visualize conserved nucleotide motifs within 24-*PHAS* transcripts. The consensus motif sequence was used as query to search candidate small RNA triggers of 24-nt pre-meiotic phasiRNAs using CD-HIT v4.8.1 (Li and Godzik, 2006) with following options: -n 8, -d 0, -g 1, -T 10 and -c 0.8. sRNAs were considered as candidate triggers when the meet following criteria: (I) 22-nt in length, (II) ≥10 raw reads and (III) an uracil as 5’-end nucleotides.

We performed target prediction for candidate sRNA triggers on pre-meiotic 24-*PHAS* precursor transcripts using sPARTA v1.26 (Kakrana et al., 2014). The sRNA-target interactions having score ≤ 4 were considered valid and used for subsequent analysis. To visualize the conservation of the motif among *PHAS* transcripts, we performed a multiple sequence alignment using MUSCLE v3.8.1551 (Edgar, 2004) with default parameters. We visualize the alignment using Jalview v2.11 (Waterhouse et al., 2009).

### Bioinformatics Analysis of RNA-seq Data

Using cutadapt v1.9.1, RNA-seq reads were pre-processed at the 3’ end using a Phred quality score ≥ 20 and trimmed reads shorter than 25-nt were discarded. Clean reads were mapped to wheat and barley reference genomes using HISAT2 v2.1.0 (Kim et al., 2015). A *de novo* reference-guide transcript assembly was performed using StringTie v2.0.1 (Pertea et al., 2015; Pertea et al., 2016) with following parameters: -c 10, -f 0.2 and -m 200. Anther-specific transcriptomes expressed in wheat and barley were then reconstructed with StringTie –merge (run with -f 0.2 and -m 200 parameters).

Transcript assemblies were compared to wheat and barley reference annotations using gffcompare utility (http://github.com/gpertea/gffcompare) and parsed in two groups of known and unknown gene/transcript loci. Unknown transcripts were analyzed in three steps. First, we annotated transcripts having homology to Rfam v14.1 sequences (Kalvari et al., 2017; Kalvari et al., 2018) using the Infernal v1.1.2 program (Nawrocki and Eddy, 2013) with default parameters. Second, transcripts without a confident match to Rfam were analyzed to annotate lncRNA by combining CREMA v1.0 (Simopoulos et al., 2018) and FEELnc v0.1.1 (Wucher et al., 2017) programs with default parameters. Third, remaining transcripts were analyzed to identify open reading frames (ORFs) using Transdecoder v5.5.0 (https://github.com/TransDecoder/TransDecoder.git). ORFs containing a CDS ≥ to 150 amino acids were aligned to the SwissProt (https://www.uniprot.org/downloads; download: 2019-01-10) protein database using the blastp option of diamond v0.9.29.130 (Buchfink et al., 2015) with following options: -e 1e-100, --matrix BLOSUM62 --more-sensitive. The final annotated assembly was used to support interpretations of sRNA regulation during anther development.

### Abundance Analyses of RNA

Prior to performing abundance analyses, we generated a read-count matrix for sRNA- and RNA-seq experiments in wheat and barley. The read-count matrix produced by ShortStack was used to perform sRNA abundance analyses. For RNA-seq assembly, reads mapping to genes of wheat and barley assemblies were counted using StringTie (run with -e and -B parameters) and a read-count matrix was extracted using a python helper script (prepDE.py) provided with StringTie. Prior to preforming the expression analysis, we filtered the read-count matrices and kept only sRNA clusters or genes with respectively ≥ 5 and ≥ 10 reads per million over a minimum of 3 samples and normalized read count abundance for library size with the TMM method by using edgeR (Robinson et al., 2010; McCarthy et al., 2012). To assess the degree of uniformity among replicates of anther at distinct developmental stages, an MDS analysis was performed with edgeR. The TMM normalized transcript count per million (TPM) was generated and used to identify co-expressed genes using the WGCNA R package (Langfelder and Horvath, 2008) to perform a weighted correlation network analysis.

### Orthology and Phylogeny Analysis of Protein Genes Involved in sRNA Pathways

To study protein-coding genes involved in sRNA pathways, we used OrthoFinder v2.3.11 (Emms and Kelly, 2015) and SonicParanoid v1.2.6 (Cosentino and Iwasaki, 2018) to perform a gene orthology inference in wheat and barley. Proteomes of three dicots (*A. thaliana, G. max* and *S. lycopersicum*) and ten monocots (*A. tauschii, B. distachyon, H. vulgare, O. sativa ssp. indica, O. sativa ssp. japonica, T. aestivum, T. turgidum ssp. dicoccoides, T. turgidum ssp. durum, T. urartu* and *Z. mays*) were included to the analysis. Additionally, unknown ORFs derived from *de novo* transcriptome assemblies in wheat and barley anther were added to these analyses. RDR, DCL, DRB and AGO reference protein sequences of *A. thaliana, Z. mays* and *O. sativa* (reported by Zhang et al., 2015 or available on UniProt database) were used to identify orthologous groups of these protein families. Retrieved protein sequences were aligned using MUSCLE v3.8.1551 (Edgar, 2004) with default parameters. Protein alignment was trimmed with trimAL v1.4.rev15 (Capella-Gutierrez et al., 2009) with the following parameter: -gappyout. Protein alignments were used to infer phylogenetic trees by maximum likelihood using IQ-TREE v1.6.12 (Minh et al., 2013; Nguyen et al., 2015; Kalyaanamoorthy et al., 2017) with following parameters: -alrt 1000 and -bb 1000. Finally, we used iTOL v4 (Letunic and Bork, 2019) to draw consensus trees.

### Data Visualization

Various plots were produced to visualize data using R programs. Circular plots were drawn using OmicCircos v.1.24.0 (Ying and Chunhua, 2015) to draw the chromosomal distribution and total abundance of phasiRNA and miRNA loci. Programs pheatmap v.1.0.10 (https://rdrr.io/cran/pheatmap/) and ggpubr v.0.2.4.999 (Wickham, 2016; Kassambara, 2018) were used to visualize abundance changes in phasiRNA through the development of anther.

### Data Availability

The complete set of raw sRNA-seq and RNA-seq reads were deposited in the Sequence Read Archive under SRA accession number PRJNA636099.

## Author Contributions

B.C.M and S.B. designed the research; S.B. performed the experiments and analyzed the data; S.P. performed miRNA target motif analysis; K.Z. supervised the microscopy experiments; S.B. and B.C.M wrote and reviewed the manuscript.

The author responsible for distribution of materials integral to the findings presented in this article in accordance with the policy described in the Instructions for Authors (www.plantphysiol.org) is: Blake C. Meyers.

## ACKNOWLEDGMENTS

We thank members of the Meyers lab, as well as Graham Moore and Azahara Martin (John Innes Centre, Norwich, UK), for helpful discussions, and Joanna Friesner for assistance with editing. We thank Mayumi Nakano for assistance with data handling. We wish to thank the Advanced Bioimaging Laboratory at the Donald Danforth Plant Science Center for support with sample preparation and imaging support. This work was supported by USDA NIFA “BTT EAGER” award 2018-09058 to B.C.M., as well as resources from the Donald Danforth Plant Science Center and the University of Missouri – Columbia. This research was enabled in part by support provided by Calcul Québec and Compute Canada.

## SUPPLEMENTARY DATA

*Supplementary Table 1. Coordinates and abundance of all miRNA detected in wheat cv. Fielder (a) and in barley cv. Golden Promise (b) and Morex (c)*.

*Supplementary Table 2. Summary of all RDR, DRB, DCL and AGO protein genes encoded by the wheat and barley genome in addition to names proposed according to their evolutionary relation to other species determinate by a phylogenic analysis*.

*Supplementary Table 3. Coordinates and abundance of all PHAS loci detected in wheat cv. Fielder (a) and in barley cv. Golden Promise (b) and Morex (c)*.

**Supplementary Figure 1.**
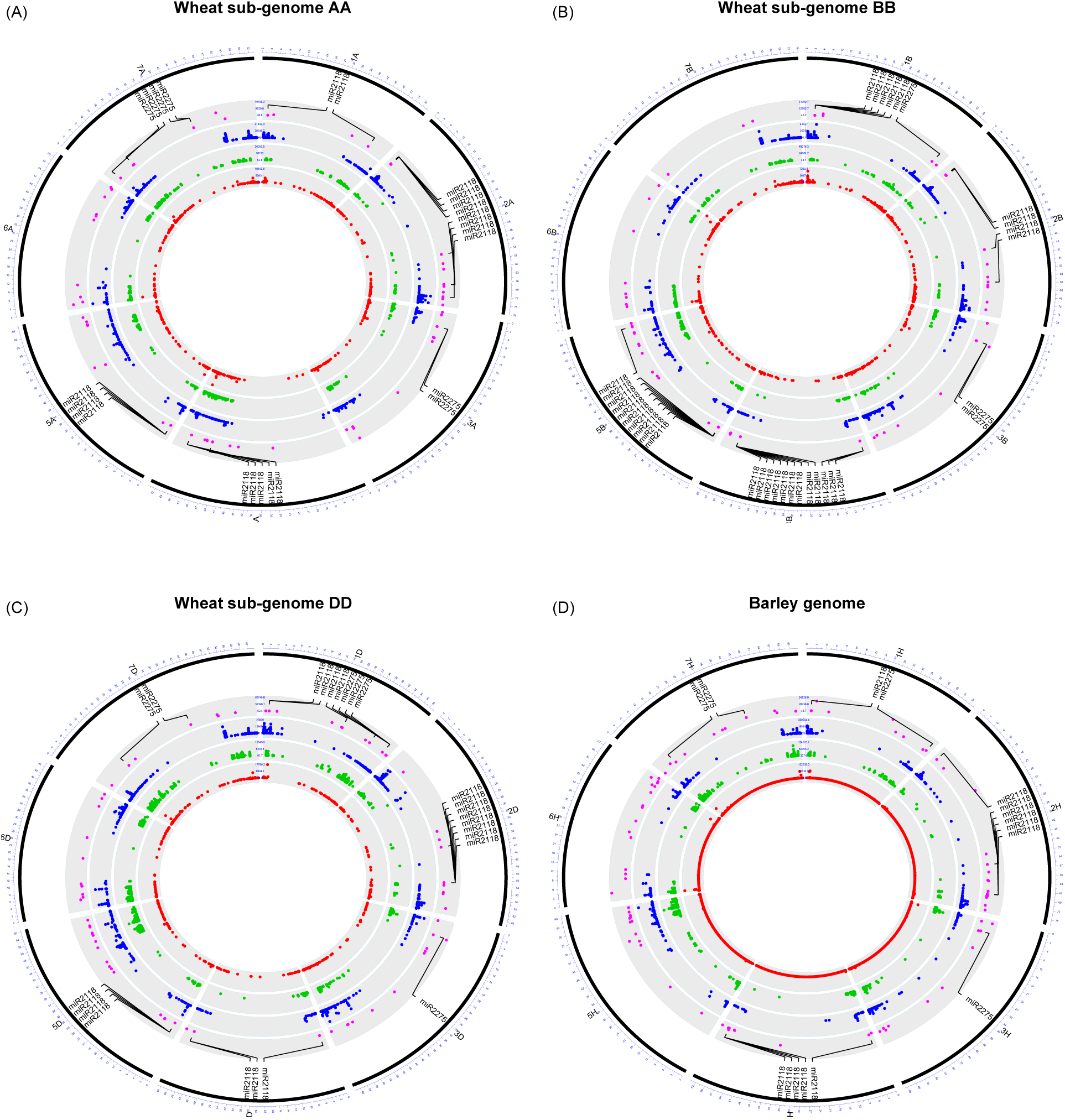
Circular plot showing the distribution and abundance of heterochromatic siRNAs (red), 21-nt (green) and 24-nt (blue) phasiRNAs as well as miRNAs (purple) annotated in wheat sub-genomes A, B, D and in barley genomes. The genomic distribution of miRNA triggers for 21-*PHAS* (miR2118) and 24-*PHAS* (miR2275) transcript cleavage are labeled.

**Supplementary Figure 2.**
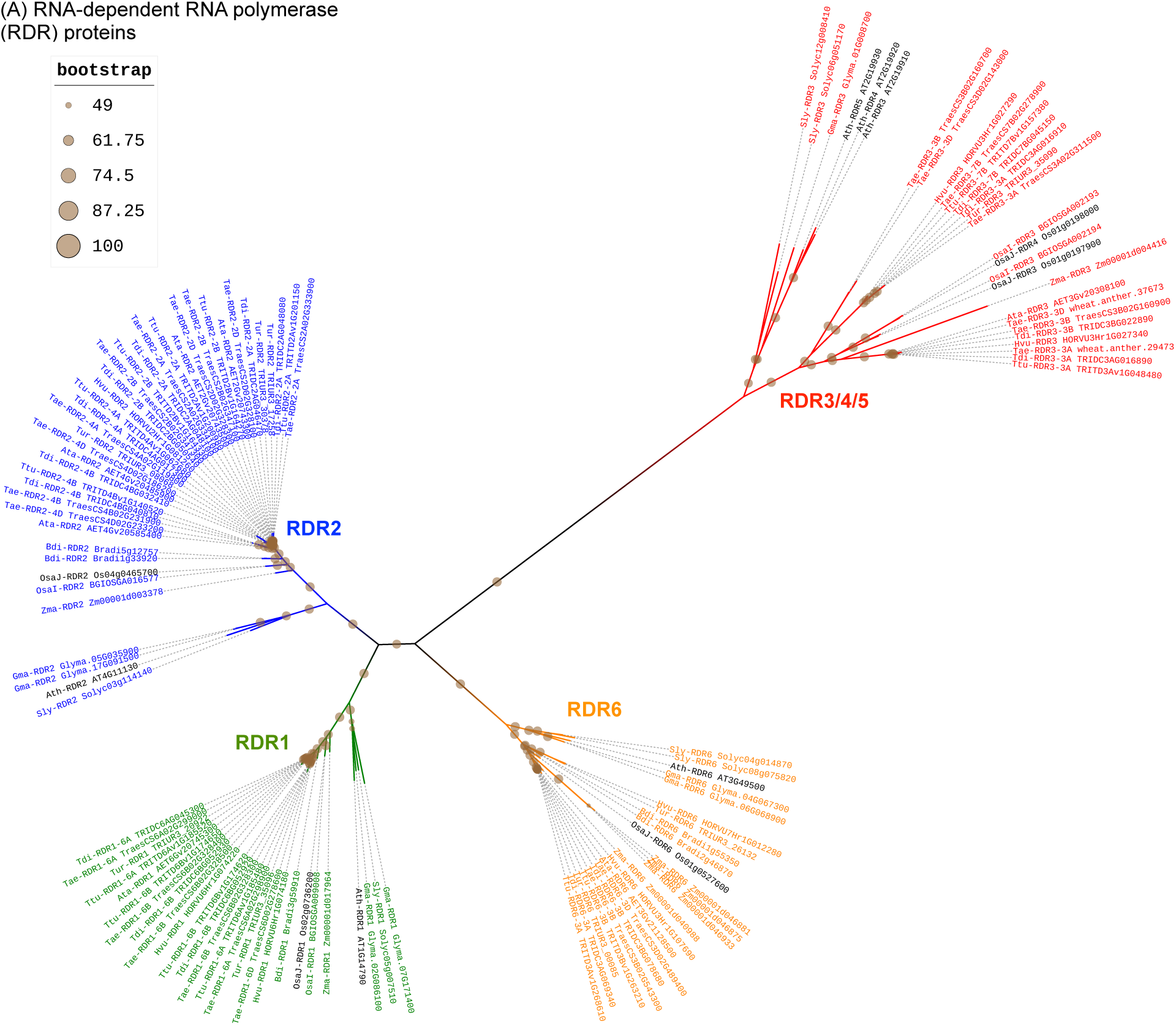
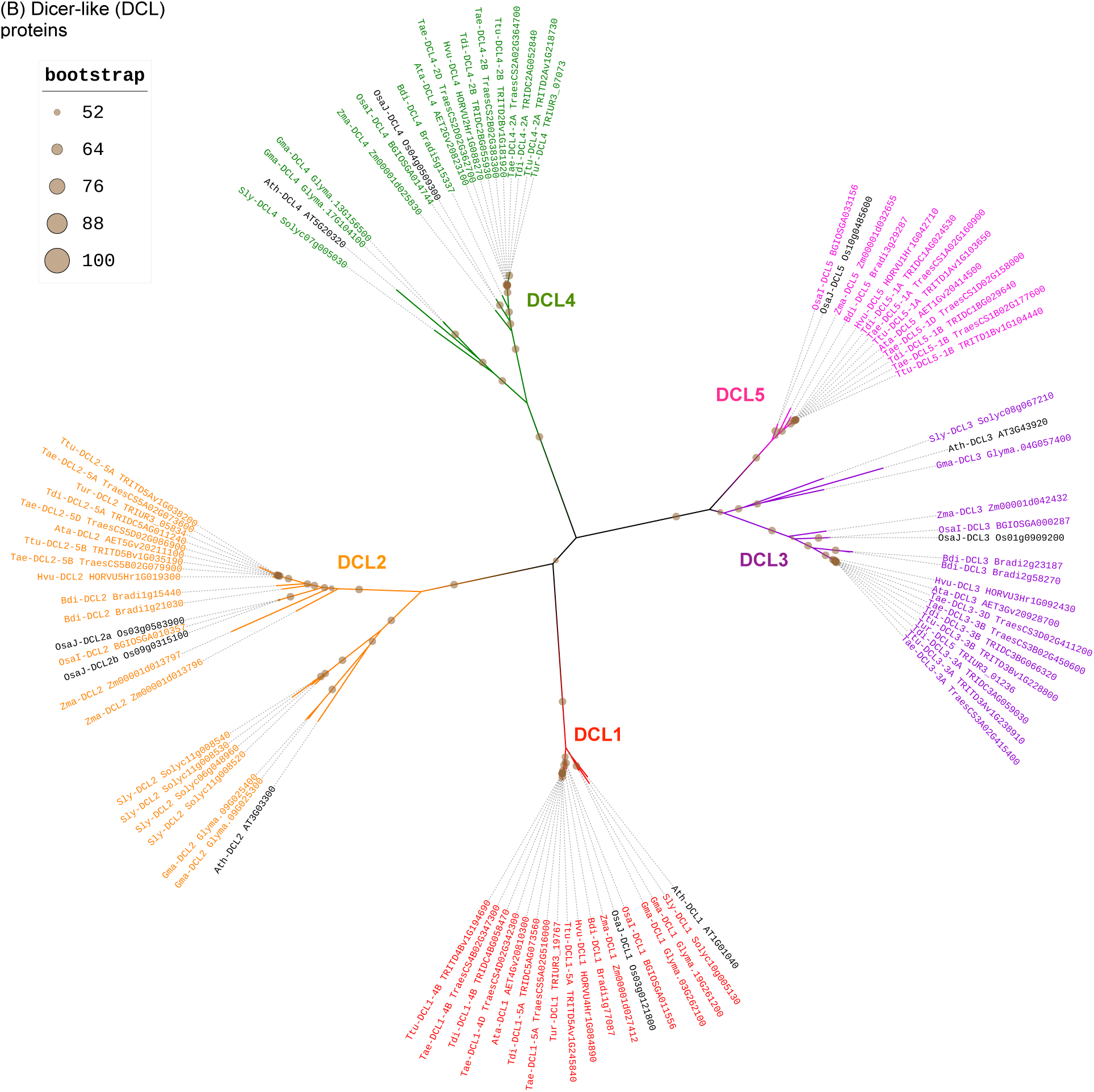
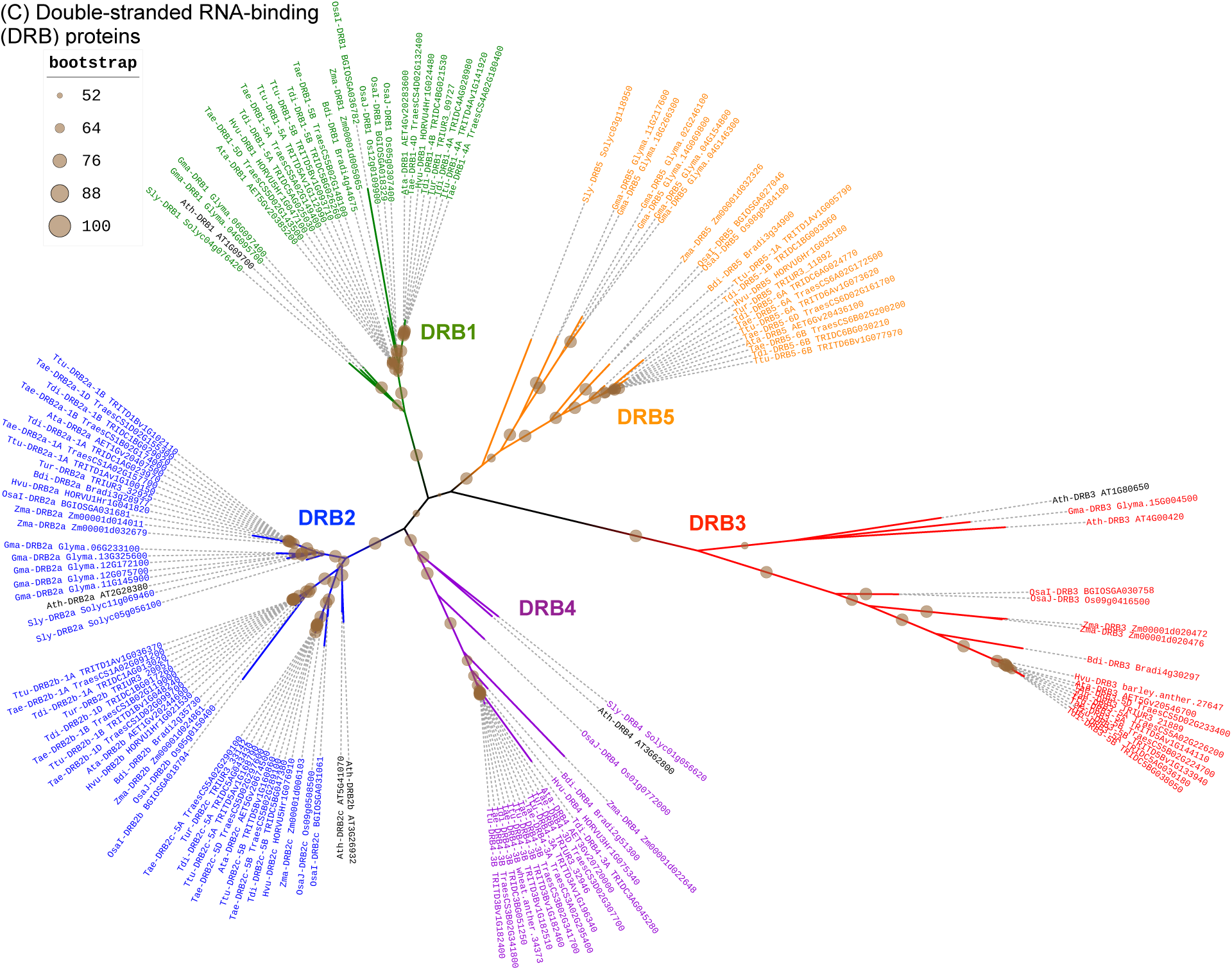
Phylogenic trees showing annotated wheat and barley orthologous genes to (A) RNA-dependent RNA polymerase (RDR), (B) Dicer-like (DCL) and (C) Double-stranded RNA-binding (DRB) proteins. Protein orthologous groups were identified using OrthoFinder and SonicParanoid. Protein alignment and phylogeny analysis were done using MUSCLE and IQ-TREE.

**Supplementary Figure 3.**
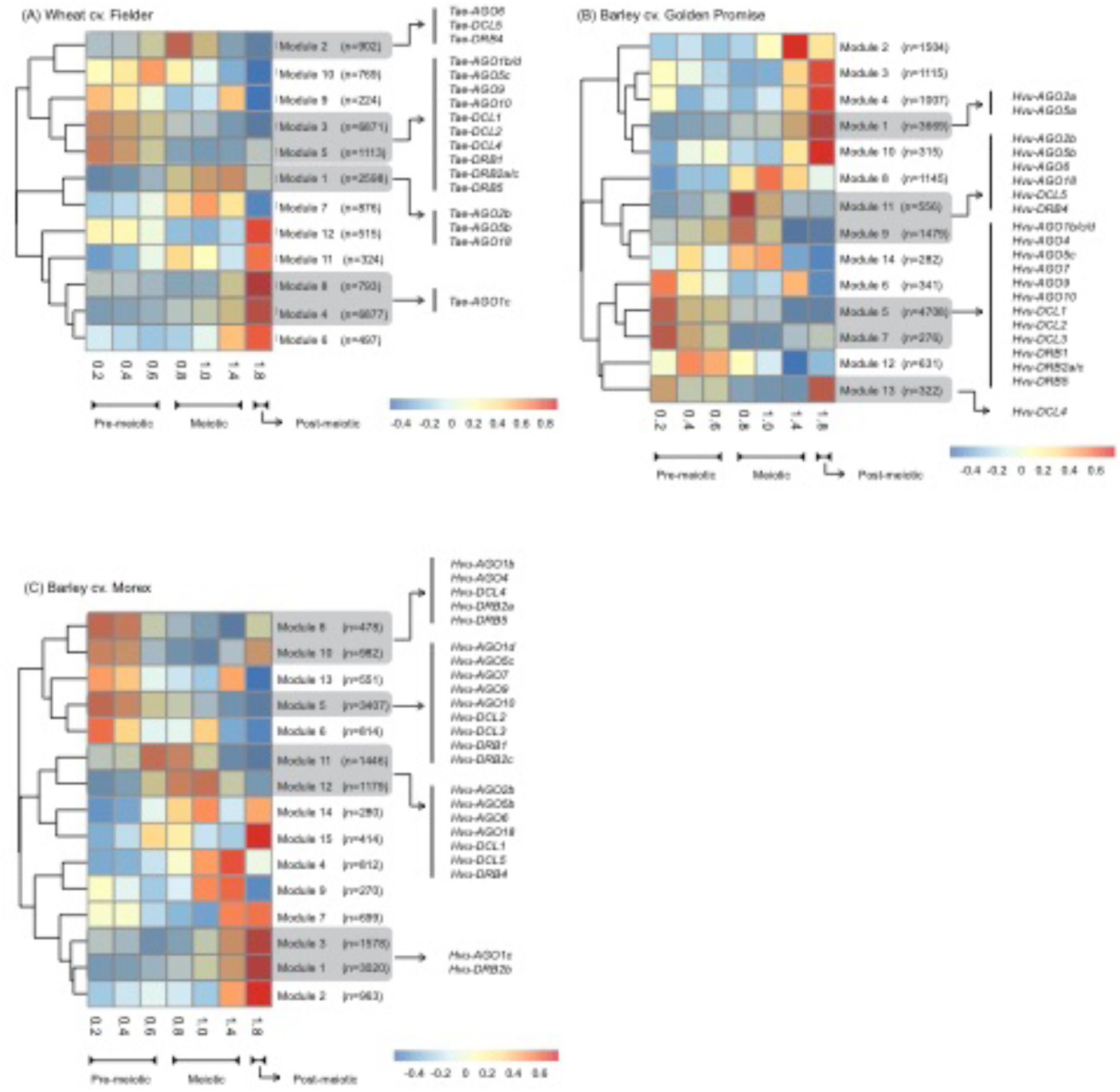
Co-expressed genes in anther of the wheat cv. Fielder (A) and barley cv. Golden Promise (B) and Morex (C). For each panel, we present a heatmap showing relative abundance of gene expression modules and *DCL, DRB* and *AGO* genes annotated in those modules. Co-expression analysis were performed using the R package WGCNA.

